# Augmenting microbial phylogenomic signal with tailored marker gene sets

**DOI:** 10.1101/2025.03.13.643052

**Authors:** Henry Secaira-Morocho, Xiaofang Jiang, Qiyun Zhu

## Abstract

Phylogenetic marker genes are traditionally selected from a fixed collection of whole genomes evenly distributed across major microbial phyla, covering only a small fraction of gene families. And yet, most microbial diversity is found in metagenome-assembled genomes that are unevenly distributed and harbor gene families that do not fit the criteria of universal orthologous genes. To address these limitations, we systematically evaluate the phylogenetic signal of gene families annotated from KEGG and EggNOG functional databases for deep microbial phylogenomics. We show that markers selected from an expanded pool of gene families and tailored to the input genomes improve the accuracy of phylogenetic trees across simulated and real-world datasets of whole genomes and metagenome-assembled genomes. The improved accuracy of trees compared to previous markers persists even when metagenome-assembled genomes lack a fraction of open reading frames. The selected markers have functional annotations related to metabolism, cellular processes, and environmental information processing, in addition to replication, translation, and transcription. We introduce TMarSel, a software tool for automated, systematic, free-from-expert opinion, and tailored marker selection that provides flexibility in the number of markers and annotation databases while remaining robust against uneven taxon sampling and incomplete genomic data.

## 1 Introduction

Phylogenetic trees serve as the cornerstone for studies ranging from estimating the age of lineages [1, 2] to comparative genomics [3] and microbial community ecology [4, 5], as they recapitulate the evolutionary history of species [6, 7]. Inference of phylogenetic trees relies on identifying phylogenetic markers from homologous sequences that descend vertically from a common ancestor (orthologs) [8–10]. In addition to orthologs, microbes harbor homologous genes that have been exchanged through horizontal gene transfer (HGT) [11, 12]. The deep divergence times of microbes, estimated at around 4 billion years [1], have entangled homologs into complex relationships that obscure the precise identification of orthologs [8, 10]. Nonetheless, new tree inference methods have been developed to bypass the identification of orthologs, thereby enabling the usage of all homologous sequences of a gene family as potential markers [13, 14]. Because the downstream applications of phylogeny heavily depend on the tree quality, it is critical to select a combination of markers that yield the most accurate tree.

Metagenome-assembled genomes (MAGs) comprise the major genomic source of microbial diversity [15]. And yet, the gold standard 16S rRNA marker used for phylogenetic surveys of microbial diversity [16] is rarely recovered from shotgun metagenomic sequences [17, 18]. Moreover, 16S rRNA-based trees reflect only the evolution of the gene rather than the set of species [19]. To improve the tree accuracy, modern phylogenetic surveys have adopted a larger number of markers involved in housekeeping functions, such as ribosomal proteins or aminoacyl-tRNA synthetases [20–24]. These markers have been selected from a fixed collection of whole genomes spanning the major microbial phyla, thereby biasing the representation of markers toward well-characterized taxa. In contrast, MAGs seldomly contain the entire genomic repertoire of a population [25, 26], and some even lack ribosomal proteins due to assembly errors [27], reducing the number of markers available for tree inference. Therefore, no one-size-fits-all set of markers exists. To account for the novel diversity and heterogeneous quality of MAGs, marker selection needs to be flexible and tailored to the input genome collection.

Although the inference of microbial trees has shifted from using a single marker [28, 29] to multiple markers [20–24, 30], their selection is restricted to universal orthologous genes, which are commonly defined as being present in 90% of genomes and existing as a single copy in at least 95% of them [31–33], severely limiting the number of markers considered. However, recent studies have shown that including gene families beyond standard universal orthologous genes leads to improvements in the accuracy of inferred trees [13, 14], suggesting the need for a comprehensive assessment of the phylogenetic signal from gene families. Here, we systematically select gene families to serve as markers for deep microbial phylogenomics tailored to the input genome collection. Leveraging recent advances in tree inference methods and genome annotation databases, we show that an expanded selection of markers yields species trees with higher accuracy than previous sets of markers in real-world datasets of whole genomes and MAGs derived from a wide range of environments. In addition to genes involved in replication, translation, and transcription, we found that markers have functional annotations related to metabolism, cellular processes, and environmental information processing, and all of them provide phylogenetic signals for tree inference. We also show that our automated, systematic, free-of-domain expertise, and tailored marker selection is robust against uneven taxon sampling and incomplete MAGs while remaining flexible in the number of markers to select and the choice of annotation database. Overall, we present a new method for Tailored Marker Selection (TMarSel), available as a software tool, that can be applied to modern genomic datasets, providing a foundation for more robust and accurate phylogenomic reconstruction.

## 2 Results

### 2.1 A vast yet unexplored gene family space for microbial phylogenomics

We survey a collection of 1,510 whole reference genomes evenly sampled across the microbial tree of life from the Web of Life 2 (WoL2) database to obtain an accurate representation of the gene family distribution in microbes. We then annotated open reading frames (ORFs) of genomes into gene families using the KEGG and EggNOG databases. KEGG gene families are scattered throughout genomes (Figure 1A), ranging from universal to lineage-specific and from single-copy to multi-copy. The traditional criteria for marker selection are restricted to genes present in at least 90% of genomes and containing one copy in at least 95% of them [31–33]. We observe that only 1% of gene families annotated from the WoL2 genomes fall within the region defined by traditional criteria. This pattern highlights the limited number of gene families used for microbial phylogenomics. The limitation is further exacerbated in gene families annotated from 793 MAGs of the Earth Microbiome Project (EMP), as MAGs do not have gene families that conform to the traditional criteria. And yet, genomes and MAGs harbor, on average, 1,289 and 846 gene families, respectively, that might add new phylogenetic signals to the tree inference process. EggNOG gene families share the same characteristics (Figure S1), suggesting that these trends are independent of the annotation database.

**Figure 1.**
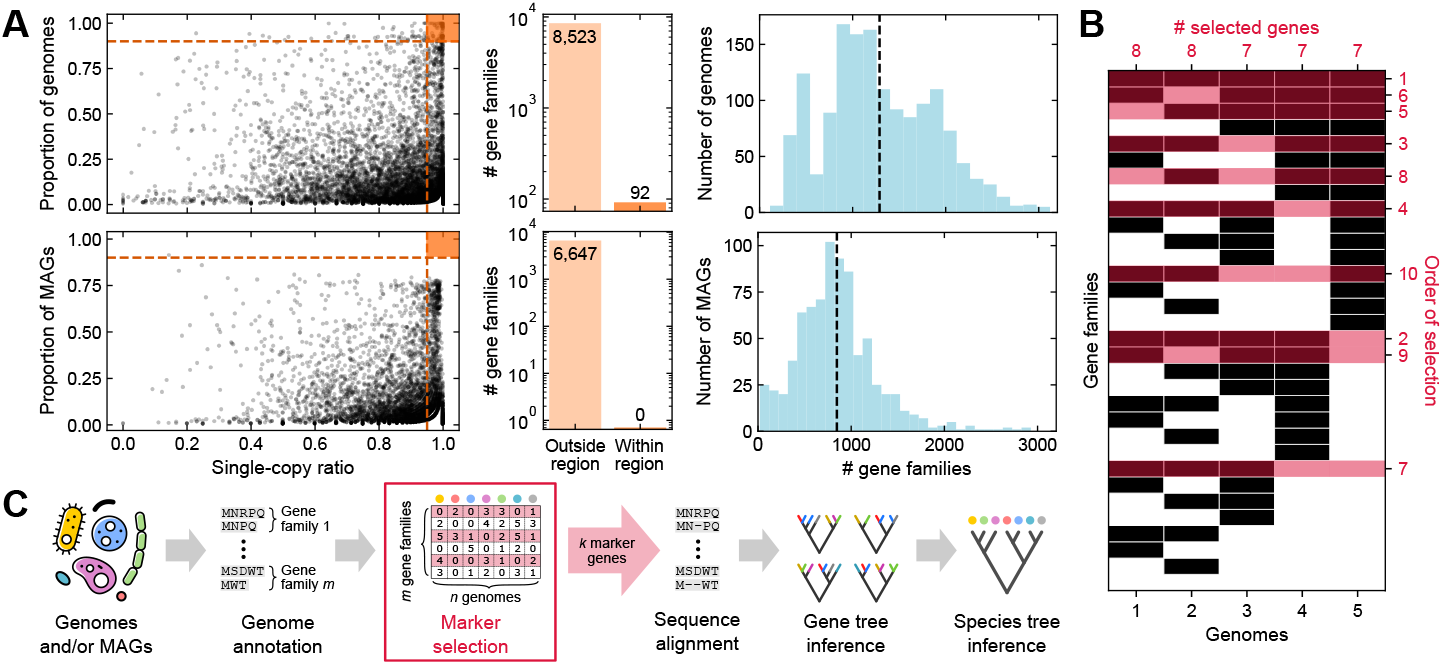
**A** Summary statistics of KEGG gene families annotated from 1,510 WoL2 genomes (top) and 793 EMP MAGs (bottom). The left panels show the gene family space defined by the proportion of genomes in which each gene family is present and the proportion of genomes in which the gene family exists as a single copy. Dashed orange lines represent traditional marker selection criteria (i.e. present in 90% of genomes and containing a single copy in at least 95% of them), and the orange region highlights the area defined by these criteria. Each dot corresponds to a gene family. The middle panels show the number of gene families outside and within the region defined by traditional criteria. The left panels show the distribution of gene families across genomes and MAGs, with dashed vertical lines indicating the arithmetic mean. **B** A simple toy case with 5 genomes and all 32 combinations of gene presence (black squares) or absence (white squares). TMarSel iteratively selects *k* = 10 markers (crimson shade) that maximize the generalized mean of the number of selected genes per species, as denoted on top. **C** Illustration of the pipeline used for species tree inference.

To assess whether these overlooked gene families can contribute phylogenetic signals to the inference process, we developed a robust yet flexible method, TMarSel, to select gene families annotated with the KEGG and EggNOG databases as markers for deep microbial phylogenomics. TMarSel employs an algorithm that iteratively selects *k* markers such that the generalized mean number of markers per genome is maximized (see Methods for details; Figure 1B). The selected markers are fed into a standard pipeline to infer species trees (Figure 1C). Briefly, for each marker, we generate a multiple sequence alignment, which is used to infer a gene tree. Because gene trees can include all the homologs of a gene family, we used the summary method ASTRAL-Pro2 [34, 35], which takes as input a set of gene trees and all their homologs, to infer a species tree. We then evaluate the quality of the inferred species trees as a direct measure of the goodness of the selected marker sets.

### 2.2 A well-balanced marker selection produces highly accurate trees in simulated data

TMarSel performs an iterative selection of markers tailored to the input genome collection. Its behavior can be customized with two parameters: the total number (*k*) of markers to select and the exponent *p* of the generalized mean. Practically, *p* biases the selection of markers toward families present in genomes with fewer gene families (if *p <* 0) or toward families present in genomes with more gene families (if *p >* 0; see Methods). To assess the impact these parameters have on the inferred trees, we first simulated toy a dataset of 50 gene families from 10 genomes across 25 replicates. In each replicate, we built a matrix containing the copy number of gene families across genomes (Figure S2A). We derived a species tree from the matrix using neighbor-joining over the Jaccard distances between genomes, upon which gene trees were also derived (see Methods). We then performed a parameter sweep for *k* and *p* and varied the maximum number of copies for each gene family and the noise present in gene trees. Noise refers to the proportion of leaves that have been randomly shuffled. We gauged the error in inferred trees as the normalized Robinson-Foulds (nRF) distance [36] between inferred and real trees, where smaller distances indicate fewer errors and vice versa. Our simulations show that selecting a large number of markers reduces the error in the species trees, primarily when noisy gene trees are used for inference (Figure 2A and standard deviations in S2B). The parameter sweep shows that *p* ≤ 0 yields the species trees with fewer errors, with *p* = 0 as an inflection point. Moreover, we observe that having multiple copies of the same marker does not improve the inference process. Instead, they negatively impact quality, as errors increase with the number of copies.

**Figure 2.**
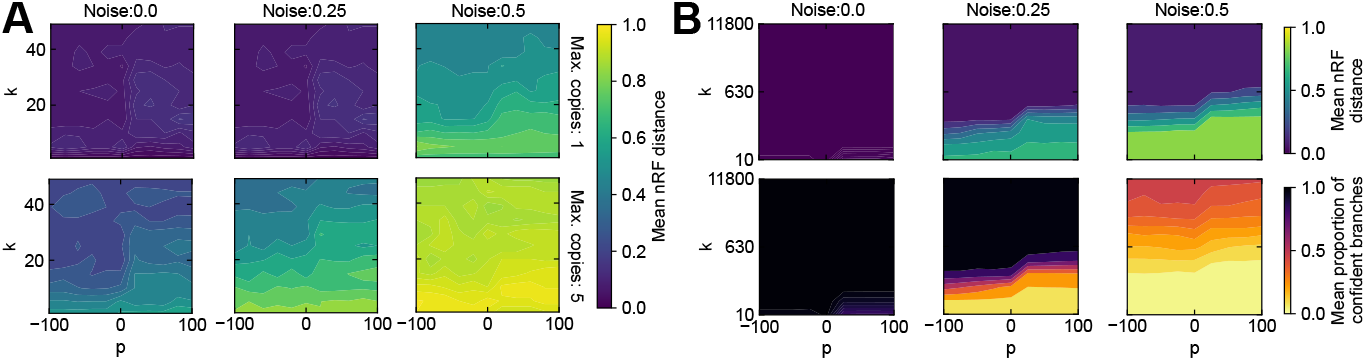
**A** Contour plots of the mean normalized Robinson-Foulds (nRF) distance between inferred and real trees from a simulated dataset of 50 gene families from 10 genomes across 25 replicates. **B** Contour plots of the mean nRF distance between inferred and real (top) and proportion of confident branches in inferred trees (bottom) from simulations of realistic prokaryotic gene families across 25 replicates. Marker selection was applied to each combination of the number of markers (*k*) and exponent (*p*). Each plot illustrates a scenario with different noise levels in gene trees (A and B) and maximum copies of gene families (only A).

We further assessed the impact of parameters on tree quality using a gene family space that resembles real datasets (Figure S2C). We simulated prokaryotic gene families using realistic duplication, transfer, and loss values [37] across 25 replicates using the phylogenetic simulator Zombi [38]. Each replicate yielded a species tree, genomes for each taxon, and gene trees for each gene family. We then built a matrix containing the copy number of gene families across taxa, introduced noise to the gene trees, and performed a parameter sweep for *k* and *p* (see Methods). In addition to the nRF, we gauged the proportion of confident branches in the inferred trees as the number of branches with a Local Posterior Probability (LPP) score higher than 0.95 out of the total number of branches, as suggested in [39]. A higher proportion is an indicator of a more accurate tree topology. As in the previous simulations, our results indicate that a large number of markers reduces the error and reinforces *p* = 0 as an inflection point that achieves the lowest error in the inferred trees (Figure 2B; standard deviations are provided in Figure S2D). The proportion of confident branches shows a similar pattern for *k* and *p* and is correlated with the error in the inferred trees. This indicates that trees with fewer errors have a higher proportion of confident branches and vice versa. The simulations also show that a saturation value occurs for *k*, at which the accuracy of inferred trees does not improve. In light of these results, we choose *p* = 0, which refers to maximizing the geometric mean number of markers per genome, for all subsequent analyses while varying *k* until reaching a plateau in quality.

### 2.3 Expanded marker selection improves the accuracy of the microbial tree of life

We next evaluated whether gene families annotated from the genomes of the WoL2 can serve as the foundational genetic elements for an accurate microbial tree of life. Because the simulation results suggested that multiple copies of the same gene family negatively impact the tree inference process (Figure 2A), we first evaluated how many copies of each gene family should be included in the analyses. Using the bit score threshold assigned during genome annotation with KEGG and EggNOG databases, we controlled the number of copies of gene families (see Methods). Our results indicate that using a low number of copies yields the tree with the highest quality as measured by the proportion of confident branches (i.e., accuracy in topology) and nRF distance to the reference phylogeny (Figure S3A). Consequently, we included only the ORFs with the highest bit score assigned to each gene family during genome annotation for marker selection.

We then benchmarked the performance of an expanded selection of markers for inferring the microbial tree of life. Figure 3A and B shows the quality of trees inferred using an increasing number of markers, ranging from *k* = 10 to 1,000. For comparison, we also included four sets of previously proposed universal markers widely adopted for deep microbial phylogenomics [1, 23, 30, 40]. Trees inferred with our markers exhibit the highest accuracy in topology and lower error to the WoL2 reference phylogeny compared to previous markers. However, the 400 universal markers from PhylophlAn3 achieve the lowest error, which is expected because the WoL2 phylogeny was reconstructed using the PhyloplhAn markers. And yet, the high quality of trees inferred from either KEGG or EggNOG gene families speaks for the robustness of TMarSel for selecting markers. Moreover, the smooth saturation curve in tree quality suggests that for any given *k*, our method can select the best set of gene families for tree inference. Filtering genomes with less than 25% of markers leads to trees with slightly lower quality across marker sets. Nonetheless, more markers per genome result in better species placement within the tree (Figure S3B). Additionally, trees inferred from all marker sets exhibit clades consistent with the GTDB taxonomy (Figure S3C), according to taxonomic accuracy metrics that measure the consistency between taxonomy and phylogeny (see Methods). Altogether, these results suggest that all marker sets recover known relationships among clades, while markers derived from KEGG and EggNOG gene families yield more accurate trees compared to previous sets.

**Figure 3.**
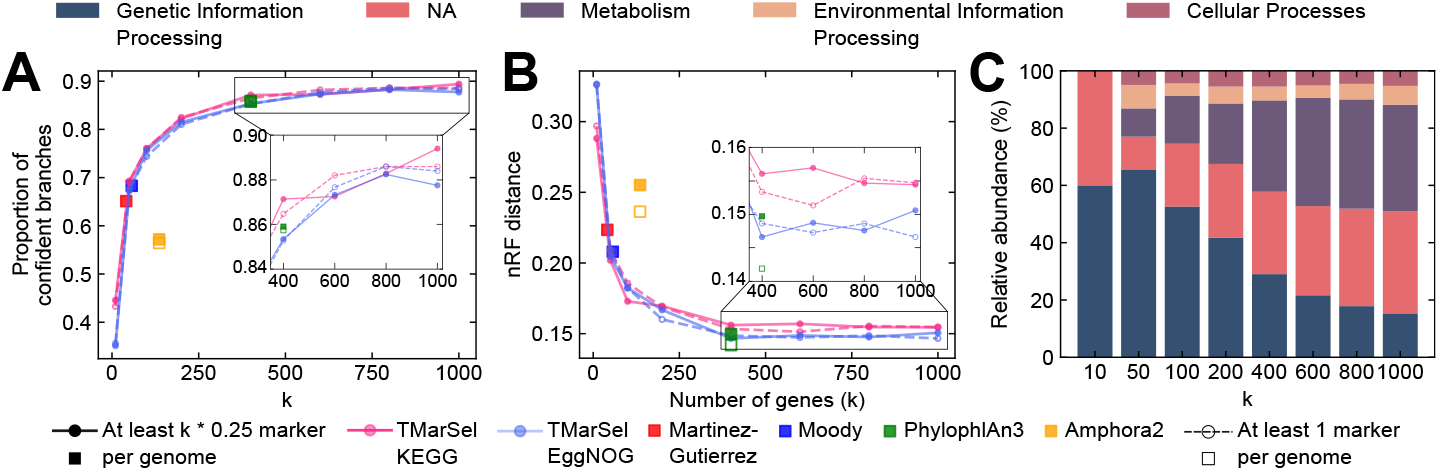
**A** Topological accuracy, measured as the proportion of confident branches, of inferred trees from 1,510 WoL2 genomes. **B** nRF distance between inferred trees and WoL2 reference phylogeny. Each dot corresponds to a tree inferred from different sets of markers (color-coded). Solid lines and filled symbols indicate trees derived from genomes containing at least 25% of markers, while dashed lines and unfilled symbols indicate trees from genomes with at least one marker. Insets are close-ups of the quality of inferred trees. **C** Relative abundance of KEGG higher functional categories (color-coded) of markers selected from KEGG gene families.

Because TMarSel iteratively selects more than twice the markers compared to the largest set available (i.e., 400 markers from PhylophlAn3), we investigated the functional classification of these new markers. Figure 3B shows the abundance of higher functional categories of markers selected from KEGG gene families. 60%, 66%, and 53% of markers have functions related to genetic information processing when 10, 50, and 100 markers are selected, suggesting that genes involved in replication, transcription, and translation are preferred. Yet, as more markers are added, the selection shifts toward gene families annotated as carrier proteins, shape-determining proteins, and others that are not assigned to higher functional categories (see Table S1 for a complete description). 38% of gene families are involved in metabolic functions, while 4% and 5% are involved in cellular processes and environmental information processing, respectively, when more than 600 markers are selected. We also observe an overlap in the functional descriptions of KEGG and EggNOG gene families (Table S1), which speaks to the robustness of selection. These results highlight the diverse functional roles of these new markers that contribute to the phylogenetic signal.

The prevalence of mobile genetic elements (MGEs) in genomes from major microbial phyla [41], suggests that genome annotations are likely to yield multiple MGE gene families. Therefore, we assessed whether gene families annotated as MGEs were selected as markers by surveying their functional description terms (see Methods). We found that putative transposases, integrases, and competence proteins were selected. Nonetheless, they account for less than 2% of markers and represent less than 13% and 6% of the total number of gene families annotated with MGE descriptions in the KEGG and EggNOG databases, respectively (Figure S4A). This showcases the robustness of TMarSel against the over-selection of MGEs. We also tested whether the taxonomic distribution of input genomes biases the selection of markers. To do this, we randomly sampled 1,000 genomes from the 15,953 WoL2 genomes across 25 replicates and counted the times each marker was selected (see Methods). We found that TMarSel consistently selects the same markers regardless of the taxonomic distribution of the input genomes (Figure S4B). This robustness suggests that our method is ideal for metagenomic datasets where taxonomic imbalance is typical.

### 2.4 Robust marker selection yields accurate trees for MAGs despite incomplete genomic data

Most of the microbial diversity comes from MAGs derived from environmental samples [15]. Microbial communities from environments as diverse as seawater, soil, sediment, or animal gut sequenced in the EMP dataset have contributed to the genomic corpus of microbial diversity [42]. To test whether marker selection with TMarSel yields accurate phylogenies for MAGs, we used 793 high-quality MAGs from the EMP dataset annotated with KEGG and EggNOG databases. Because no reference phylogeny exists for the EMP dataset, we evaluated the quality of the inferred trees using only the proportion of confident branches, which measures the accuracy of topology. Similar to the WoL2 results, we found that using the ORFs with the highest score for each marker results in more accurate topologies (Figure S5A). Despite the overall quality of trees being lower than those from the WoL dataset, the TMarSel markers selected from KEGG and EggNOG gene families produce trees with the highest accuracy compared to previous sets of universal genes (Figure 4A). The increase in accuracy is more pronounced when more markers are used in the inference process (*k* ≥ 400), though a plateau is reached at *k* = 800. Moreover, filtering MAGs with less than 25% of marker yields trees with ∼ 5% more confident branches. Although the filtering step decreases the number of MAGs across all marker sets, trees inferred with TMarSel retain more MAGs than the next-best set of universal genes from PhylophlAn3 (Figure S5B). The functional annotation and proportion of MGEs in TMarSel markers are congruent with those selected from the WoL2 dataset (Figure S5C and D).

**Figure 4.**
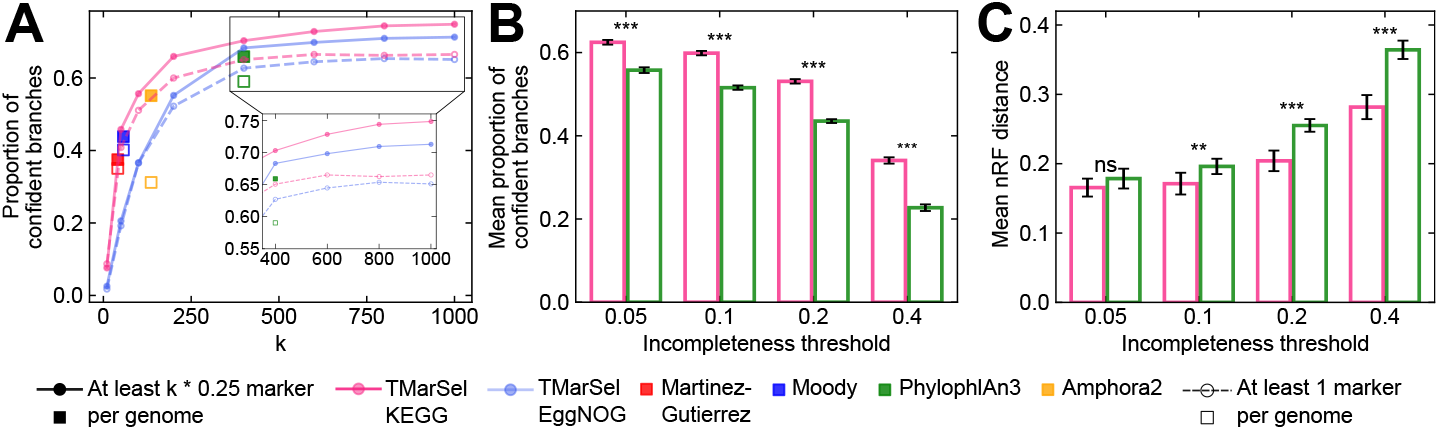
**A** Topological accuracy, measured as the proportion of confident branches, of trees inferred from 793 complete MAGs from the EMP dataset. Inset is a close-up of the quality of inferred trees. Symbols indicate trees inferred from different marker sets (color-coded), while line style and filled or unfilled symbols distinguish trees inferred from MAGs containing at least 25% of markers versus those containing only one marker. **B** Mean topological accuracy of trees inferred from simulations of increasingly 793 incomplete MAGs. **C** mean nRF distance between trees inferred from incomplete MAGs to complete MAGs. The incompleteness threshold refers to the proportion of ORFs removed from each MAG. Unfilled bars indicate that MAGs contained at least one marker. Error bars indicate standard deviations and asterisk the significance level of Mann-Whitney tests between the quality of trees inferred from KEGG versus Phylophlan markers. Significance levels: ns: p-value *>* 0.05; *: p-value ≤ 0.05; **: p-value ≤ 0.01; ***: p-value ≤ 0.001.

MAGs represent draft microbial genomes, but most MAGs do not capture all the genomic content of a microbe [25]. To assess whether accurate trees can still be inferred, we simulated increasingly incomplete MAGs across 10 replicates. The incompleteness threshold is defined as the proportion of ORFs removed from a MAG (see Methods). We then selected markers (*k* = 400) from KEGG-derived gene families, as they produced the most accurate trees. To obtain a comprehensive picture of the impact of incompleteness, MAGs with at least one marker were included. We then measured the accuracy in the topology of trees inferred from incomplete MAGs and the mean nRF distance (error) between trees inferred from complete MAGs and incomplete MAGs. We also included trees inferred with the 400 universal genes from PhylophlAn3 for comparison and performed Mann-Whitney tests with multiple testing corrections to assess whether differences in tree quality were significant. As expected, the accuracy of the trees decreases with the incompleteness threshold because fewer gene families are available for selection. Nonetheless, on average, trees inferred from TMarSel markers have 8% more confident branches than trees inferred from PhylophlAn3 universal genes across incompleteness thresholds (Figure 4B). In addition, the robustness of our markers is evident in the low errors of inferred trees, which increase slower at higher thresholds compared to the trees inferred from PhylophlAn3 markers (Figure 4C). Overall, these results suggest that a tailored marker selection from MAGs yields more accurate trees compared to previous sets of markers, even when MAGs lack a fraction of ORFs.

## 3 Discussion

In this study, we assessed the phylogenetic signal of gene families for microbial phylogenomics using TMarSel, a robust yet flexible method for marker selection. We show that an expanded and tailored selection of markers can improve the accuracy of phylogenetic trees across simulated and real-world datasets of whole genomes as well as incomplete MAGs.

TMarSel provides a systematic exploration of the gene family space because it expands the source of markers to gene families instead of only orthologs. Among the currently available methods that model the evolutionary histories of gene families for tree inference, only ASTRAL-pro2 scales efficiently with a large number of genomes and genes [14], allowing us to assess the impact of different parameter combinations on the quality of the species trees. Although ASTRAL-pro2 only accounts for gene duplication and loss, its quartet-based approach is robust against HGT [43]. Moreover, the taxonomic consistency of clades, from the phylum to the genus levels, in trees inferred from TMarSel markers speaks for the robustness of our results despite the pervasiveness of HGT among microbes. Because we select gene families as markers, TMarSel is also compatible with other tree inference methods that actively account for gene duplication, loss, and transfer [44–46].

While previous marker sets were selected from a pre-defined collection of sequences and had a fixed number of markers, we offer more flexibility in both aspects. First, we rely on functional databases to obtain gene families from the input genomes, upon which TMarSel is applied. Our results show that markers selected from either KEGG or EggNOG yield trees with similar accuracy. This suggests a potential generalization to alternative databases, such as PFAM, UniRef, and MetaCyc. The choice of the database may depend on the focus of the study and downstream applications. For instance, KEGG links gene families to metabolic pathways, chemical reactions, enzymes, and other high-level functions [47–49]. Whereas EggNOG, in addition to functional annotations, provides evolutionary details of gene families [50, 51]. We can leverage the continuous updates of these databases to provide a state-of-the-art set of markers. Second, rather than setting a fixed number of markers, TMarSel can select the best *k* markers despite uneven taxon sampling (Figure S4B). We show that increasing the number of markers yields more accurate trees. The increase in accuracy follows a saturation curve that suggests an optimal range for *k*, which can be identified by selecting an increasing number of markers until reaching a plateau. Since our selection method follows an iterative approach, smaller sets are distilled versions of larger marker sets. The iterative nature of TMarSel also suggests that the first selected markers contribute more phylogenetic signal, as shown in the saturation curves of tree quality (Figures 3 and 4). Therefore, if computational resources are limited, a small set of markers still yields accurate trees. It has been suggested that at least 30 markers should be used for microbial phylogenomics [52]. However, the minimum number depends on the input data, as 50 markers yield trees with different proportions of confident branches when genomes or MAGs are used. As a result, the minimum number of markers increases when inferring phylogenies from MAGs.

MAGs comprise a large portion of the prokaryotic diversity, and their representation in genomic databases will only grow as more environmental samples are sequenced [15, 53]. However, the integration of MAGs into phylogenomic pipelines comes with challenges. First, as we have shown, MAGs do not have gene families that conform to the traditional criteria for selection. Second, even high-quality MAGs, gauged as complete by CheckM, do not contain the entire genomic diversity of a microbial population [25, 26], and some MAGs even lack ribosomal RNA and ribosomal protein genes due to assembly errors [27]. TMarSel effectively addresses the first challenge, as it can systematically explore the gene family space and select the best *k* markers tailored to the input set of MAGs. The second challenge is more complex because incomplete MAGs inherently lack genomic data. And yet, we show that it is still possible to select markers with strong phylogenetic signals, yielding trees with high accuracy while allowing for partial recovery of the tree topology. Although current annotation databases cannot assign all the ORFs from MAGs to gene families, recent advances in functional annotation can assign the missing ORFs to novel metagenome gene families [54], which TMarSel can leverage to find new markers for microbial dark matter.

TMarSel effectively selects the optimal combination of markers from gene families that serve as the foundational genetic elements for inferring accurate phylogenetic trees from a tailored input of whole genomes and MAGs despite uneven taxon sampling and incomplete genomic data. Since TMarSel primarily relies on the presence-absence patterns of gene families, it remains agnostic to the taxonomy of the input genomes or MAGs, as well as functional constraints, allowing for the selection of markers with diverse functional roles. We also show that marker selection can be agnostic to evolutionary rate, alignment quality, and non-vertical evolution. However, further evaluations of these properties are desirable to identify rogue markers that are detrimental to the inference process. In addition, we have only assessed the performance of our method for taxa spanning across the microbial tree of life. And yet, the robustness of our results indicates a potential generalization to fine-grained taxonomic groups, as well as other annotation databases and tree inference methods.

## 4 Materials and Methods

### 4.1 Marker gene selection

We represented gene families *U* = {*u*_1_, …, *u*_*m*_} across genomes *V* = {*v*_1_, …, *v*_*n*_} as a 2D matrix *A*_*m*×*n*_ where entries are positive if gene family *u*_*i*_ exists in genome *v* _*j*_ and its values correspond to the number of times the gene is identified in the genome (i.e., copy number). Gene families not existing in a given genome were represented with a zero in *A*. To select a set of *k* marker genes *G* = {*g*_1_, …, *g*_*k*_}, *k < m*, we devised an algorithm that, in each *k* iteration, selects the gene *g* that maximizes the objective function: argmax 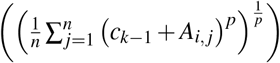, where *c*_1× *n*_ is a cumulative vector containing the copy number of genes already selected that were removed from *A* (see Algorithm 1). Note that our cost function is analogous to the generalized mean with exponent *p*. Small values of *p* shift the cost function toward small values, selecting genes present in genomes with fewer gene families. In contrast, large values of *p* will select genes from genomes with many genes. Because *A* contains zeroes, our cost function will return zero for *p* ≤ 0 (Figure S2A). To avoid this issue, we added a pseudocount of 0.1 to *A* when selecting marker genes for all values of *p*.

#### Algorithm 1

Algorithm for marker selection

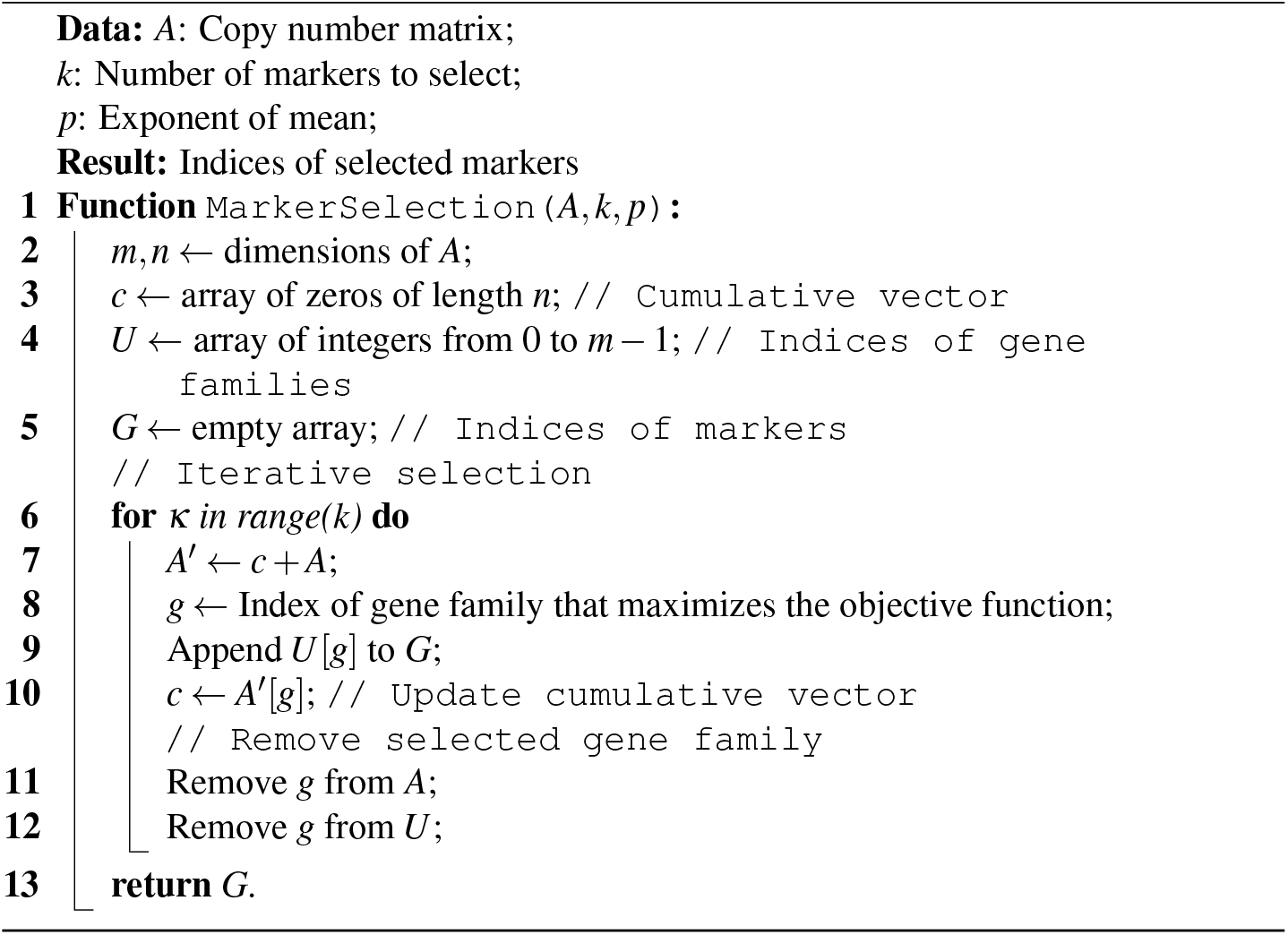

### 4.2 Toy simulations

To assess the impact of parameters *p* and *k*, we simulated multiple copy number matrices as follows: i) Fix the number of gene families (*m* = 50) and genomes (*n* = 10). ii) For each genome, randomly draw a number between zero and one from a uniform distribution to represent the fraction of gene families in the genome. For instance, a value close to zero corresponds to a genome with few gene families and vice versa. iii) For each genome, generate a presence-absence vector of gene families, where a value of one (i.e., presence) is sampled with a probability equal to the fraction of gene families from step ii. In contrast, a value of zero (i.e., absence) is sampled with a probability of 1 - fraction of gene families. Thus, step iii produces a presence-absence matrix of gene families across genomes. iv) To add copy numbers over the presence-absence matrix from step iii, we sampled a number between one and a maximum number of copies, set beforehand, for each gene present within a gene family. All steps were implemented using NumPy v1.26.0 [55] while fixing a seed for reproducibility. This procedure was repeated for 25 replicates, yielding 25 different copy number matrices later used to select marker genes. For each replicate, we then performed a parameter sweep for 11 values of *k* and *p* ranging from *k* = 1 to *k* = *m* − 1 and from *p* = − 100 to *p* = 100 (Figure S2A).

To generate a species tree from the copy number matrix, we first calculated the Jaccard distance for each pair of genomes to obtain a distance matrix. We then applied neighbor-joining implemented in Scikit-bio v0.6.2 over the distance matrix to generate a species tree. To obtain gene trees, we took the species tree as a template, and for each gene family, we removed the genomes that did not contain the gene family. Hence, at this stage, gene trees contain only a single copy of each genome. We then added gene copies as sister branches of a given genome. Thus, for example, if a gene family is present in four out of ten genomes with copy numbers ranging from one to five (e.g., 1, 3, 5, 2), the resulting gene tree will have 11 leaves in total from four genomes, representing all the copies that stem from duplication or transfer events. Because we do not infer gene trees from sequences, we introduced noise into gene trees to simulate uncertainties that may arise during multiple sequence alignment and impact the gene tree inference process. Specifically, noise here refers to the proportion of branches that are randomly shuffled. Scikit-bio v0.6.2 was used to manipulate trees.

### 4.3 Realistic simulations

Because the toy simulations are unrealistic, we simulated realistic prokaryotic gene families using Zombi [38]. We first generated a species tree under a birth-death model for 500 units of time with speciation and extinction rates per unit of time set to 0.04 and 0.03 per unit of time, as suggested by Louca et al. [56]. We then simulated genomes along the branches of the species tree using 10,755 and 5,577 bacterial and archaeal gene family-wise values of duplication (D), transfer (T), and loss (L) benchmarked in [37] while leaving the rest of the parameters as default. At the end of the simulation, Zombi outputs the real species tree and the genomes of each taxon along with real gene trees of each gene family. From the genomes, we built the copy number matrix that was later used to select marker genes. We repeated the simulations for 25 replicates while fixing a seed for reproducibility. For each replicate, we then performed a parameter sweep for 11 values of *k* and *p* ranging from *k* = 1 to *k* = *m* − 1 and from *p* = − 100 to *p* = 100. We also introduced noise to the gene trees to simulate uncertainties that may arise during multiple sequence alignment.

### 4.4 Web of Life 2 and Earth Microbiome Project datasets

The Web of Life (WoL2) contains 15,953 prokaryotic species with a single representative high-quality genome and taxonomic annotations from the Genome Taxonomy Database (GTDB) R207, as well as a reference phylogeny [57]. The WoL2 dataset is publicly available at https://ftp.microbio.me/pub/wol2/. Due to the burden of computational resources, we selected complete reference genomes from the WoL2, ensuring that each taxonomic group, from phylum to family, had at least 10 taxa. This resulted in 1,510 genomes spanning the entire microbial diversity that will be used for genome annotation and marker selection.

The Earth Microbiome Project (EMP) [42] contains 811 high-quality Metagenome Assembled Genomes (MAGs) from 32 environments across the globe and are publicly available at https://www.globus.org/, endpoint emp 500 public. After genome annotation (see below), we inspected whether the number of ORFs matched the number of gene families. We discarded 18 poorly annotated MAGs with a low number of gene families compared to the number of open reading frames (ORFs). This resulted in 793 MAGs that will be used for all subsequent analyses.

### 4.5 Genome annotation

To generate the copy number matrix for marker gene selection, we need to annotate the genomes and MAGs. First, we used Prodigal v2.6.3 [58], in single-genome mode with the genetic code table specified according to taxonomy, to predict ORFs for genomes of the WoL2. ORFs from MAGs of the EMP dataset were predicted with Prokka v1.14.6 [59]. We then used the KEGG Orthology release 102.0+ [49] and evolutionary genealogy of genes: Non-supervised Orthologous Groups (EggNOG) v5.0 [50] databases to annotate the ORFs into gene families with KOfam-Scan and EggNOG-mapper v2 [60], respectively. In the KEGG annotation, we selected only prokaryotic KEGG Orthologs (KOs) with an e-value lower than the threshold defined in the database [61], thus minimizing false positive assignments. In the EggNOG annotation, we only selected the assignments at the broadest taxonomic level since we are interested in inferring a tree for diverse microbial species. The annotation of ORFs into gene families was then used to generate a copy number matrix *A*_*m*×*n*_ with *m* gene families and *n* genomes.

We used the hierarchical classification from KEGG to map KOs to higher functional categories. For this, we counted the number of times a given gene is mapped onto a functional category. The counts in each category were then normalized by the total number of counts and multiplied by 100 to obtain a relative abundance. EggNOG only provides a functional description for each gene family. Therefore, we could not map its gene families to higher functional categories.

In addition, we identified Mobile Genetic Elements from KEGG and EggNOG gene families using the following description terms: baseplate, capsid, excisionase, DUF4102, pf00665, KilA-N, ORF11CD3, phage, portal, tail, terminase, tape, T5orf172, viral, virion, conjugal, conjugation, conjugative, DotD, IV secretory, IV secretion, MobA, mobilisation, mobilization, MobL, DUF955, plasmid, relaxase, TcpE, TraG, TraL, TraM, DDE, pf01609, IS66, IstB, transposase, transposon, transposition, anti-restriction, antirestriction, integrase, integration, K02238, K02242, K02243, K02244, K02245, K02246, K12296, K04096, K06198, K07343, as suggested in [62].

### 4.6 Controlling copy numbers

Since ORFs mapped to gene families by KOfamScan or EggNOG contain summary statistics, we focused on the bit score value to control the number of copies of a gene family present within a genome. We used the bit score rather than the e-value, as the former is independent of the database size. For every genome, we identified the maximum bit score assigned to each gene family. We then excluded ORFs that had a bit score below a certain threshold that represents proximity to the maximum bit score. For example, a threshold of one will only include the best hits of each gene family per genome. In contrast, a zero threshold will include all the ORFs. Thus, the threshold represents the number of gene copies of each gene family per genome to include for marker selection.

### 4.7 Impact of uneven taxon sampling on marker selection

To assess whether an uneven taxon sampling of the input set of genomes biases the selection of markers, we randomly sampled 1,000 genomes from the 15,953 WoL2 genomes across 25 replicates. Because the parameter *p* shifts the cost function toward smaller (if *p <* 0) values, the selection of markers is biased toward gene families from genomes with fewer gene families. Thus, this parameter directly impacts the selected markers, and the input set of genomes further amplifies its impact. However, it can be adjusted to find an optimal value for *p* that yields the same markers even when the taxonomic distribution in the input genomes is uneven. Therefore, in each replicate, we selected *k* = 400 markers from the 1,000 genomes while varying the parameter *p* of the cost function from *p* = 0 to *p* = − 10, as the simulations have shown that negative values yield accurate trees.

We then counted how many times each marker was selected across replicates. To obtain a more quantitative measure of the robustness against taxonomic imbalance, we calculated the Jaccard index of selected markers across replicates for each *p* value. A higher index indicates that the same markers are selected regardless of the taxonomic distribution of the input set of genomes, whereas a lower index suggests that different markers are selected in each replicate.

### 4.8 Pipeline for species tree inference

Once markers have been selected, we retrieved all the ORF sequences associated with them. Depending on the experimental condition, we excluded those genomes with less than *k* × 0.25 marker genes. Protein sequences of each marker gene were then aligned using UPP2 [63] with default parameters. UPP2 is a multiple-sequence aligner designed for large datasets containing sequence length heterogeneity that might arise under large insertion or deletion events or due to incomplete assembly [64, 65]. In the first stage, UPP2 selects a set of full-length sequences and computes a backbone alignment and an unrooted tree. A collection of profile Hidden Markov Models (pHMM) is then built for each subset of sequences in the backbone tree, and the remaining sequences are inserted into the backbone alignment. We chose the optimal number of backbone sequences using the backbone query split algorithm from UPP2. Data pipeline errors, such as sequencing, assembly, genome annotation, or alignment errors, substantially impact phylogenetic reconstruction as they increase the noise in the dataset. We used TAPER [66] with default parameters to remove such errors within alignments. TAPER is an outlier section algorithm that removes amino acids based on a divergence score computed along genomic positions and species [66].

We used the clean alignments for Maximum Likelihood tree reconstruction with FastTree v2.1 [67] using the Lee-Gascuel (LG) model of amino acid substitution, as a previous study showed that the LG model best explains substitutions in a majority of prokaryotic marker genes [57]. This step produced a set of *k* gene trees. We used TreeShrink [68] to remove pipeline errors that have escaped detection at the sequence level and resurfaced as suspiciously long branches in the phylogeny. TreeShrink removes leaves that inflate the tree diameter, defined as the maximum distance between any two leaves in the tree [68]. In our case, a leaf corresponds to a protein sequence of a gene within a genome.

We then used ASTRAL-Pro2 [35] to estimate a species tree from the set of gene trees. ASTRAL-Pro2 combines information from gene trees to maximize a measure of quartet similarity between them and the species tree [34]. The quartet similarity measure is defined as the fraction of partitions between all combinations of four species, with a common ancestor originating from a speciation event that shares the same branching structure (i.e., topology) as the real species tree under gene duplication and loss scenarios [34]. Because only speciation events count toward the measure, ASTRAL-Pro2 identifies orthologs from gene trees containing all the homologs of a gene family [34, 35]. This flexibility is ideal because the genome annotation step produces gene families that contain orthologs, paralogs, and xenologs.

### 4.9 Sets of previous markers

To benchmark the usefulness of our marker genes for inferring phylogenetic trees, we compiled four datasets of marker genes previously proposed and used for deep microbial phylogenomics. i) 41 single-copy universal marker genes tested for their phylogenetic signal in multidomain phylogenetic reconstruction [23]. ii) 57 single-copy universal markers, which were used to estimate the age of the Last Universal Common Ancestor [1]. In the case of the 41 universal markers from Martinez-Gutierrez, the pHMMs of each gene were already provided, whereas, for the 57 universal markers from Moody, we built the pHMMs from the multiple sequence alignments with hmmbuild from HMMER v3.4 [69]. We then used hmmsearch to search for homologs of the markers in the genomes of the WoL2 and MAGs from the EMP dataset. For each gene, only the ORF with the top bit score was extracted and used in the pipeline for species tree inference. Other marker sets we benchmarked are: iii) 136 universal markers from AMPHORA2 [30] available in PhyloPhlAn3 [40], and iv) 400 single-copy universal markers first proposed in [24, 70] and part of PhyloPhlAn3. To identify homologs of these two last sets of universal genes on the WoL2 and EMP datasets, we ran PhyloPhlAn3 v3.1.68 with high diversity and fast parameters. We then took the identified ORFs and used them in the pipeline for species tree inference. This approach ensures that species trees from all marker genes were inferred using the same pipeline, thus guaranteeing a fair comparison.

### 4.10 Quality of species trees

To evaluate the quality of species trees inferred with our marker genes, i) we calculated the Robinson-Foulds distance between the inferred tree and the WoL2 reference phylogenetic. The RF distance counts the different number of ways to divide a set of taxa by removing a branch [36]. We further normalized the RF distance by the sum of internal branches between the two trees to obtain a number between zero and one, where zero indicates two identical trees and vice versa. DendroPy v4.6.1 [71] was used for all tree distance calculations. ii) We also measured the quality of the inferred trees using the Local Posterior Probability (LPP) scores from ASTRAL-Pro2, which is a measure of confidence of each branch based on gene tree quartet frequencies. An LPP higher than 0.95 was used to classify a branch as highly confident, as suggested in [39].

iii) We gauged the taxonomic consistency of clades in a phylogenetic tree using the taxonomic accuracy metrics from [70] and the standard microbial GTDB taxonomy R207. Taxonomic precision captures the notion that phylogenetically closer species must share a common taxonomic label. The precision of a clade is calculated as 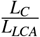 where *L*_*C*_ is the sum of branch lengths of the members of the clade (*C*) and *L*_LCA_ is the sum of branch lengths of all the descendants of the Lowest Common Ancestor (*LCA*) of the clade. Thus, if all the members of the clade form a monophyletic subtree, the precision is one (highest possible). If any member of the clade is scattered, the denominator grows faster than the numerator, thereby reducing the precision. Taxonomic recall, on the other hand, captures whether taxonomically similar taxa are grouped close in the phylogeny. The recall of a clade is calculated as 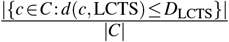, where *d* is the distance between a member of the clade *c* and the Longest Consistent Taxonomic Subtree (LCTS) and *D* is the diameter (i.e., the branch length separating the most distant leaves) of the LCTS. The recall, therefore, calculates the number of taxa outside the LCTS. The LCTS is defined as the internal node with the largest number of children that are part of the clade and are monophyletic themselves.

iv) In addition, we assessed the placement of each taxon from an inferred tree relative to the reference tree. We first calculated a distance matrix from the number of edges (i.e., internal branches) connecting every pair of taxa in the inferred and reference trees. Since each row in the distance matrix represents a vector containing the distance of a taxon to the other taxa in the tree, we calculated the cosine distance between the vectors of the same taxa in inferred and reference trees. Thus, if a taxon has the same placement in the inferred tree as in the reference tree, the distance vectors have the same direction, resulting in a cosine distance of zero. We expect well-placed taxa to have lower cosine distances to the reference tree. To assess the overall trend of placement, we applied a least squares regression implemented in SciPy v1.11.3 [72].

### 4.11 Simulations of incomplete MAGs

To simulate genome incompleteness, we randomly sampled contigs and sequentially selected ORFs in each contig until satisfying an incompleteness threshold, defined as the proportion of ORFs to be removed. The threshold is specified as a proportion of the total number of ORFs in the MAG. For instance, a threshold of 0.1 in a MAG containing 1,000 ORFs will result in 100 ORFs removed. Once all the ORFs to be removed had been selected, we removed them from the genome annotation files that relate ORFs to gene families produced by KOfamScan and EggNOG mapper. We then used the remaining gene families to build the copy number matrix for marker gene selection. For each incompleteness threshold, we repeated the simulation for ten replicates while fixing a seed for reproducibility. In each replicate, a different contig and, subsequently, ORFs were sampled. Thus, our simulations yield MAGs with different genomic compositions.

For each replicate, we selected markers (*k* = 400) from the incomplete set of MAGs and inferred a species tree. We also inferred trees using the 400 universal markers from PhylophlAn3 for comparison. The trees were then evaluated for the proportion of confident branches. In addition, we calculated the nRF between trees inferred from incomplete MAGs and trees inferred from complete MAGs. To assess whether the differences in quality between our markers and PhylophlAn markers were statistically significant, we performed Mann-Whitney tests and corrected the p-values with the Benjamini-Hochberg method implemented in SciPy v1.11.3.

## Author contributions

Conceptualization, H.S.-M. and Q.Z.; investigation, H.S.-M. and Q.Z.; formal analyses, H.S.-M. and Q.Z.; methodology, H.S.-M. and Q.Z.; software, H.S.-M. and Q.Z.; data curation, H.S.-M. and Q.Z.; writing – original draft, H.S.-M.; writing – review & editing, H.S.-M., Q.Z., and X.J.; visualization, H.S.-M., Q.Z., and X.J.; supervision, Q.Z. and X.J.; funding acquisition, Q.Z. and X.J.

## 5 Data and code availability

The datasets generated and analyzed, as well as the code used to produce the results presented in this manuscript, are publicly available on GitHub (https://github.com/HSecaira/AugmentingPhyloSignalMicrobes) and Zenodo (https://doi.org/10.5281/zenodo.14991373), under the BSD 3-Clause license. The source code of TMarSel is hosted on GitHub (https://github.com/HSecaira/TMarSel/tree/main), together with documentation and test data sets.

## Acknowledgments

We thank Dr. Siavash Mirarab for valuable suggestions on the selection method and tree inference pipeline. We thank Dr. Justin Shaffer for organizing and sharing the EMP dataset. We are grateful to Matthew Aton for sharing computational resources to run analyses and to members of the Jiang lab for their insightful feedback on the manuscript. This work utilized the computational resources of the Sol Supercomputer at Arizona State University [73].

## Supplementary Material

**Supplementary Table S1.** Functional descriptions of 1,000 markers selected from KEGG and EggNOG gene families. KEGG description also contains the pathways and higher functional category of each marker. Available as an Excel spreadsheet.

**Supplementary Figure S1.**
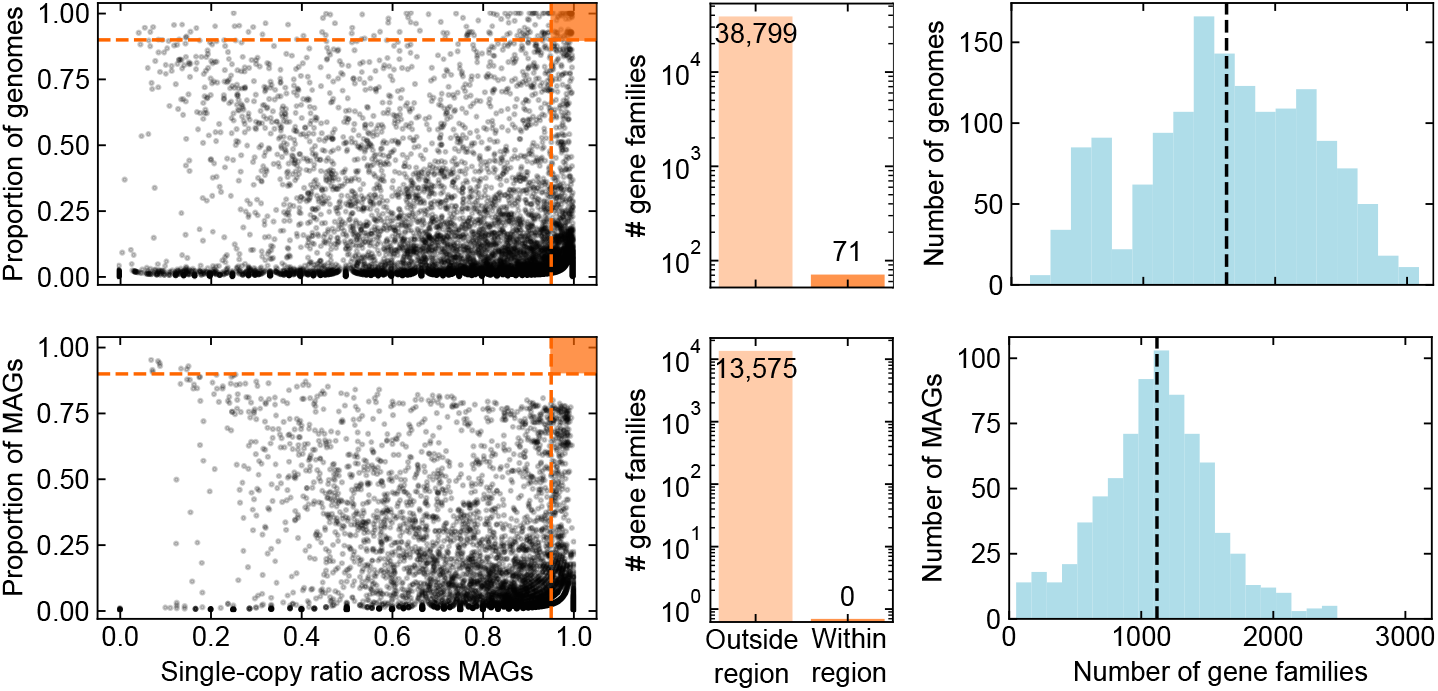
Summary statistics of EggNOG gene families annotated from WoL2 genomes (top) and EMP MAGs (bottom). The left panels show the gene family space defined by the proportion of genomes and MAGs in which each gene family is present and the proportion of genomes and MAGs in which the gene family exists as a single copy. Dashed red lines represent traditional marker selection criteria (i.e. present in 90% of genomes and containing a single copy in at least 95% of them), and the red region highlights the area defined by these criteria. Each dot corresponds to a gene family. The middle panels show the number of gene families outside and within the region defined by traditional criteria. The left panels show the distribution of gene families across genomes and MAGs, with dashed vertical lines indicating the mean.

**Supplementary Figure S2.**
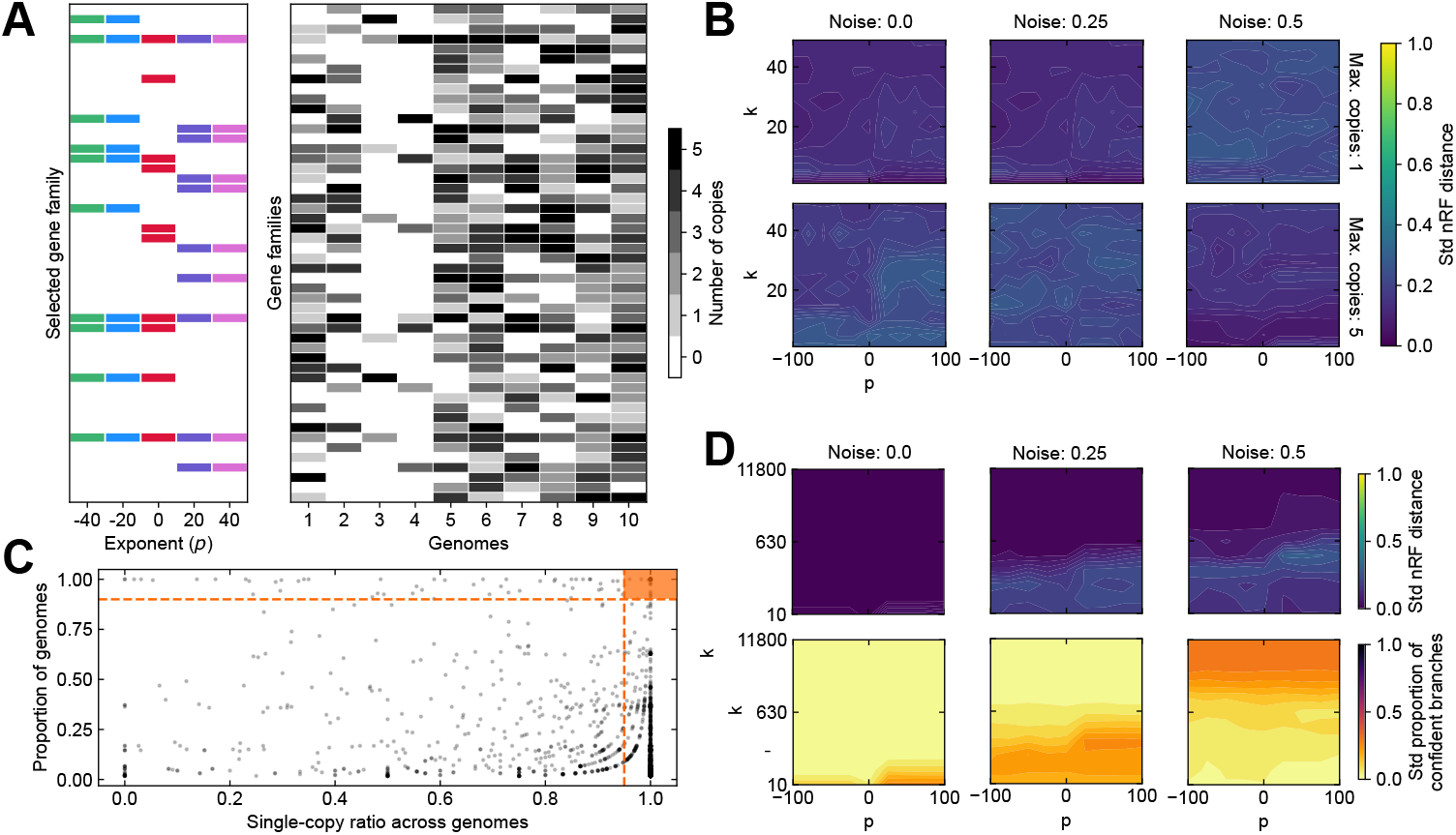
**A** Simulated toy dataset of 50 gene families from 10 genomes, and up to 5 copies of each gene family per genome. The probability of a gene’s presence per species was randomly sampled between 0 and 1, thereby leaving some genomes (such as genomes 3 and 4) sparse in gene presence. A series of exponent (p) values of the cost function were tested to select markers. As shown, higher p values select genes from genomes with many gene families, whereas lower p values tend to select genes from genomes with fewer gene families (notably genome #3). **B** Standard deviations of normalized Robinson-Foulds distance from toy simulations. **C** Gene family space of the prokaryotic gene families simulated using realistic duplication, transfer, and loss values. Dashed orange lines represent traditional marker selection criteria, and the orange region highlights the area defined by these criteria. Each dot corresponds to a gene family. **D** Standard deviations of the proportion of confident branches from realistic simulations.

**Supplementary Figure S3.**
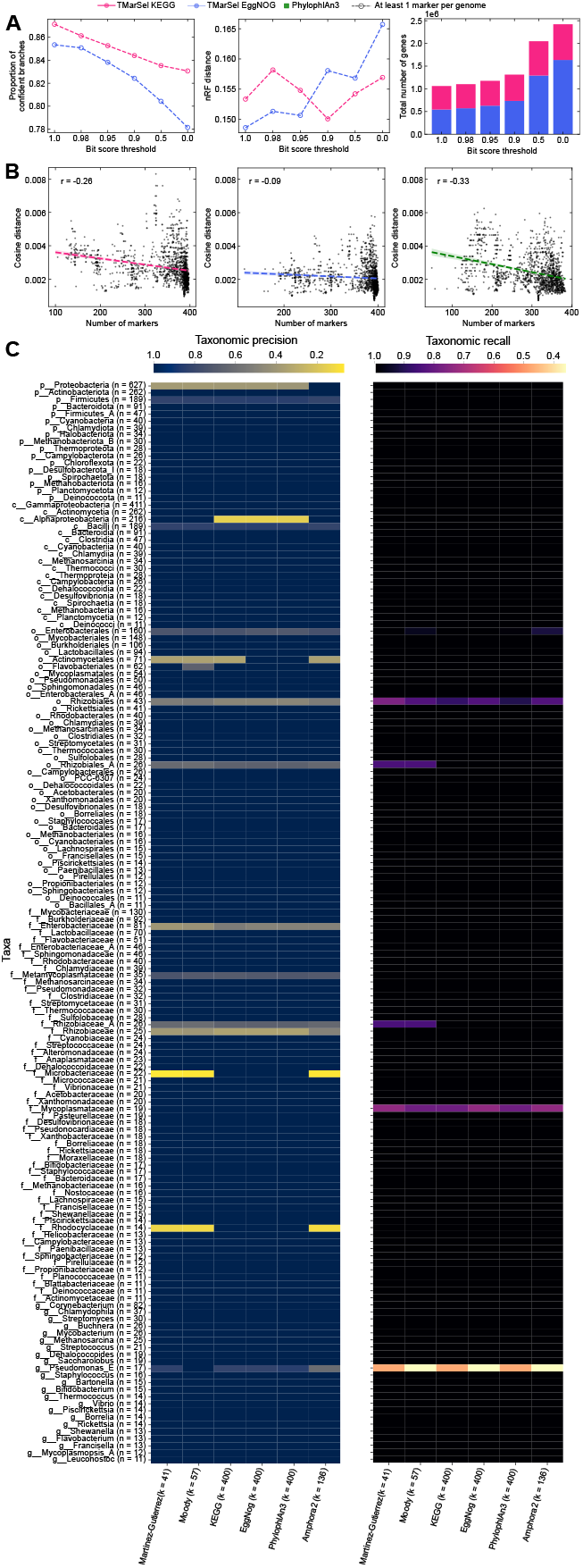
**A** Impact of copy number on the quality of trees inferred from KEGG and EggNOG gene families (color-coded) derived from the WoL2 dataset. The quality was measured as the proportion of confident branches (left panel) and normalized Robinson-Foulds distance (middle panel). The bit score threshold represents the number of copies included for each of the 400 selected markers used in tree inference (right panel). Dashed lines and unfilled symbols indicate that trees were inferred from genomes containing at least one marker. **B** Placement of genomes from inferred trees relative to the WoL2 reference phylogeny. Each dot corresponds to the cosine distance between the vectors spanned by a given genome in the inferred tree and the reference phylogeny. Dashed lines indicate the least squares regressions and *r* indicates the Pearson correlation coefficient. Genomes with more markers have a better placement in the inferred tree relative to the reference phylogeny. **C** Taxonomic accuracy metrics for each taxonomic group (rows) across marker gene sets (columns). Higher values indicate greater consistency between taxonomy and phylogeny.

**Supplementary Figure S4.**
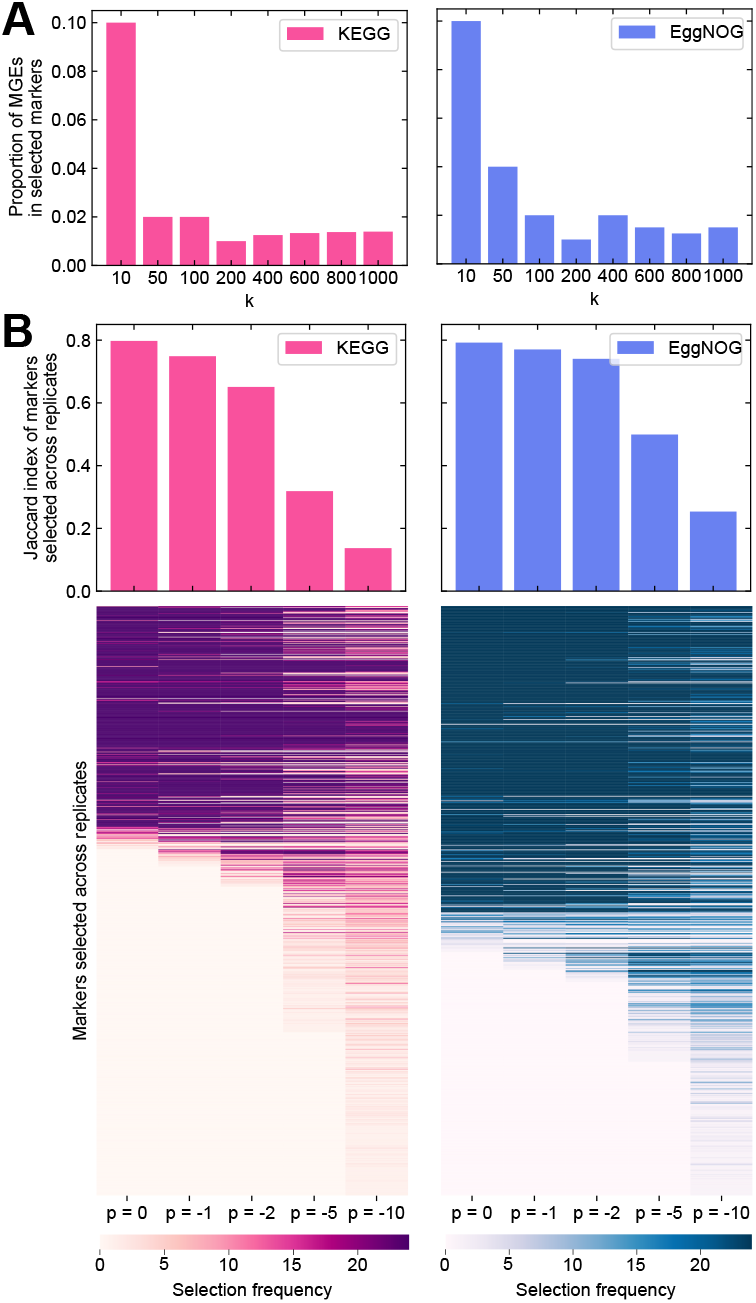
**A** Proportion of Mobile Genetic Elements in markers selected from KEGG and EggNOG gene families. Each bar corresponds to a different number of markers (*k*). **B** Robustness against uneven taxon sampling. Heatmaps show the number of times markers (rows) were selected using different values of the *p* parameter (columns) across 25 replicates. Barplots indicate the Jaccard Index of selected markers across 25 replicates for each value of the *p* parameter. A higher index indicates the same markers are selected regardless of the taxonomic distribution of the input set of genomes. Markers are consistently selected when *p* = 0. KEGG and EggNOG markers are color-coded.

**Supplementary Figure S5.**
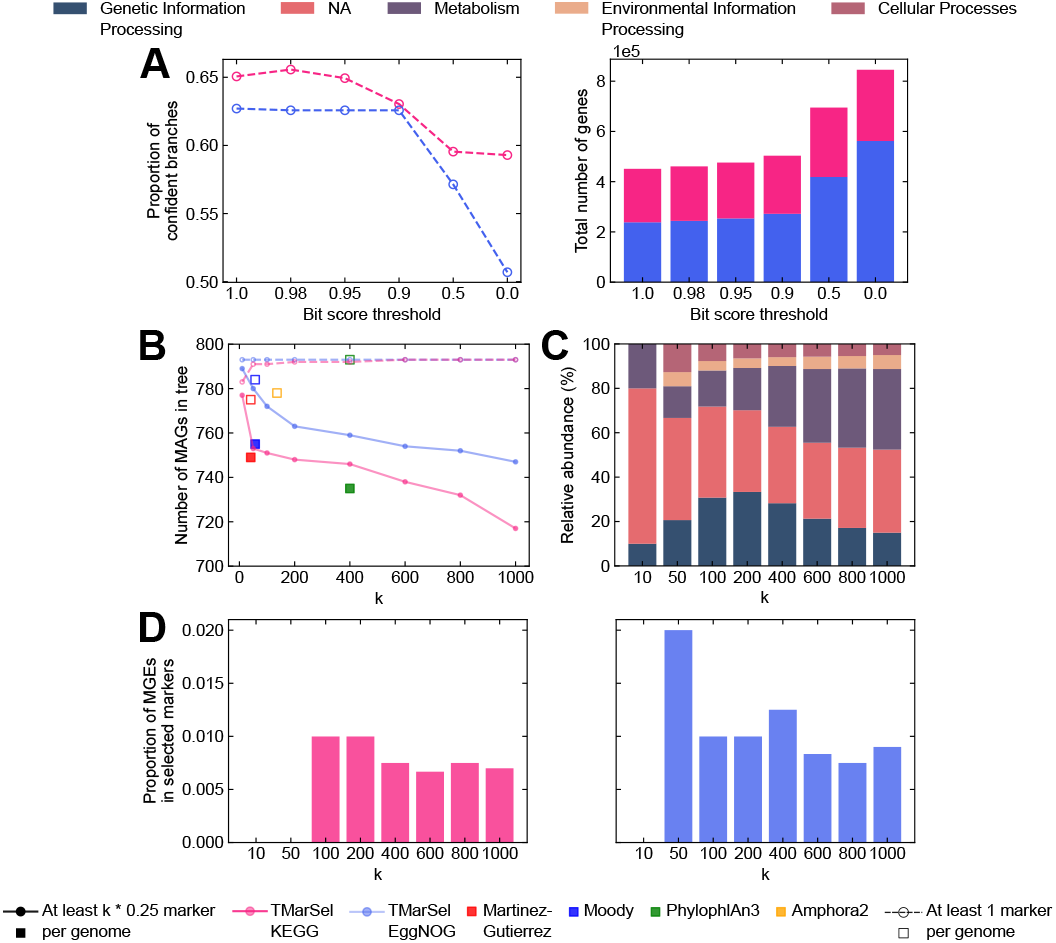
**A** Impact of copy number on the quality of trees inferred from KEGG and EggNOG gene families (color-coded) derived from the EMP dataset. The quality was measured as the proportion of confident branches (left panel) and normalized Robinson-Foulds distance (middle panel). The bit score threshold represents the number of copies included for each of the 400 selected markers used in tree inference (right panel). **B** Number of MAGs across trees inferred from different marker sets (color-coded). Dashed lines and unfilled symbols indicate that trees were inferred from MAGs containing at least one marker. **C** Relative abundance of KEGG higher functional categories (color-coded) of markers selected from KEGG gene families derived from MAGs. **D** Proportion of Mobile Genetic Elements in markers selected from KEGG and EggNOG families. Each bar corresponds to a different number of markers (*k*).

